# Short-term fasting of a single amino acid extends lifespan

**DOI:** 10.1101/2023.08.10.552880

**Authors:** Tahlia L. Fulton, Mia R. Wansbrough, Christen K. Mirth, Matthew D. W. Piper

**Author notes:** Corresponding author: Matthew D. W. Piper. Co-first authors. Co-senior authors.

## Abstract

Diet and health are strongly linked, though the strict changes in diet required to improve health outcomes are usually difficult to sustain. We sought to understand whether short term bouts of amino acid-specific modifications to the diet of *Drosophila melanogaster* could mimic the lifespan and stress resistance benefits of dietary restriction, without the requirement for drastic reductions in food intake. We found that flies fed diets lacking the essential amino acid isoleucine, but otherwise nutritionally complete, exhibited enhanced nicotine tolerance, indicating elevated detoxification capacity. The protection from isoleucine deprivation increased with the duration of exposure, up to a maximum at 7-day isoleucine deprivation for flies 2, 3, or 4 weeks of age, and a 5-day deprivation when flies were 5 weeks of age. Because of these beneficial effects on toxin resistance, we intermittently deprived flies of isoleucine during the first 6 weeks of adulthood and monitored the effect on lifespan. Lifespan was significantly extended when flies experienced short term isoleucine deprivation at 3 and 5 weeks of age, regardless of whether they were also deprived at 1 week. These results indicate that short-term bouts of isoleucine deprivation can extend lifespan and highlight its cumulative and time-dependent benefits. Interestingly, we found that isoleucine-deprived flies lost their protection against nicotine within three days of returning to fully fed conditions. Therefore, the mechanisms underlying lifespan extension may involve transient damage clearance during the bouts of isoleucine deprivation rather than sustained enhanced detoxification capacity. These data highlight a new time-restricted, nutritionally precise method to extend life in *Drosophila melanogaster* and point to a more manageable dietary method to combat ageing.

## Introduction

Dietary restriction (DR), which refers to reduced nutrient intake without causing malnutrition, has been repeatedly shown to extend lifespan^1,2^. There are several ways to implement DR to increase lifespan. One way is to manipulate dietary intake for the duration of life, such as caloric restriction^3–5^, or by altering the nutritional composition of food, such as by decreasing the protein to carbohydrate ratio^6^. Another way is to manipulate the diet transiently, such as intermittent fasting^7^. Each of these methods is associated with compliance issues because of our evolved appetites. Since the mechanisms for lifespan extension are complex and unresolved, it remains unknown if each type of DR extends lifespan in the same way. If these pathways converged, we would expect that transient bouts of nutrient-specific manipulation should enhance lifespan. Defining such a precise protocol would open the opportunity for more targeted investigations of DR’s mechanisms and uncover a highly feasible path to implementation.

Lifespan extension is not the only benefit that can come from DR. Temporary bouts of nutrient restriction can cross protect mice against a model of surgical stress^8–10^ and flies against ingested toxins^11^. Cross-protection, or hormesis, has previously been proposed as a strategy to combat ageing^12^. Mitohormesis is a popular theory which suggests that low level mitochondrial ROS production triggers protective responses that ultimately guard cells from accumulating molecular damage^13^. Previously, we have shown that short-term single amino acid deprivation protects flies (*Drosophila melanogaster*) against the neurotoxin nicotine^11^. This protection requires the amino acid sensor GCN2 (General Control Non-derepressible 2) and can be mimicked by pharmacological suppression of the central growth regulator, mTORC1 (mechanistic Target Of Rapamycin Complex 1). Together, these results suggest a mechanism whereby amino acid deprivation activates GCN2 and suppresses mTORC1, which triggers detoxification pathways that can metabolise nicotine. Enhanced detoxification capacity has been implicated in lifespan extension, as one of the evolutionarily conserved signatures of long-lived insulin mutants is enhanced expression of the detoxification system^14^. For example, when detox pathways are elevated in *Caenorhabditis elegans*, by manipulating their transcriptional regulator Nrf2 (Nuclear factor erythroid 2-related factor 2), lifespan is extended^15^. If enhanced detoxification capacity increases the clearance of molecular damage that causes ageing, then perhaps lifespan can be extended by short bouts of amino acid deprivation.

To test this, we subjected flies to short bouts of isoleucine deprivation and assessed the effect on lifespan. Our previous work showed us that the benefits of amino acid deprivation are highly context dependent, therefore we first assessed how isoleucine deprivation could enhance nicotine resistance at various ages. We then used these findings to design schedules of short-term isoleucine deprivation and test their effects on lifespan.

## Methods

### Fly husbandry

Experiments were conducted using adult female, white-eyed *Drosophila melanogaster* of the Dahomey strain (wDah). Outbred wDah stocks are maintained with overlapping generations in a high-density population cage of approximately 10,000 individuals on a sugar yeast (SY) diet^16^ (Table S1). Experimental flies were reared from egg to adult on a SY diet at a controlled density^17^ and to standardise mating status, newly emerged adult flies were kept in mixed cohorts on fresh SY diet for 2 days following eclosion. Flies were then lightly anaesthetised with CO_2_ and female wDah were sorted into vials containing a complete synthetic diet^18^ (Table S2) in cohorts of 5 flies per vial. The complete synthetic diet contained an exome matched ratio of amino acids^19^, and an isoleucine dropout was prepared in the same way except isoleucine was omitted^11^. Media were prepared in advance and stored for up to 4 weeks at 4°C. Fly survival was recorded using the software DLife^17^. For flies exposed to media alone, survival was recorded when flies were transferred to fresh media every Monday, Wednesday, and Friday. For flies exposed to nicotine, survival was recorded 3 times a day at 7am, 1pm and 7pm every day until all flies were dead. Experiments and stocks were maintained at 25°C, 60% humidity and a 12:12 hour light:dark cycle.

### Nicotine-laced medium

Free base nicotine (1000mg/mL; [Sigma Aldrich: N3876]) was diluted in absolute ethanol to a concentration of 25mg/mL. The nicotine-laced medium was prepared by aliquoting 100µL of diluted nicotine solution into a vial containing 3mL of cooled, gelled complete synthetic medium (final concentration of 0.83mg/mL nicotine in vials). Vials were then kept in a fabric cover at room temperature for 24-48h to ensure an even dispersion of nicotine and that the ethanol had evaporated. We have previously shown that vials laced with ethanol alone had no effect on survival of flies during this assay period^11^. Nicotine-laced food was prepared as needed, 48h in advance of use, and was not stored.

### Figures and data analysis

All analyses were completed using R^20^ (version 4.2.2) and R Studio^21^ (version 1.4.1106) and we created all plots using ggplot2^22^. All data and scripts are publicly available at: [To be made freely available through Figshare].

To determine whether the independent variables of a model could explain variation in the data, we initially analysed the models using a type II or III ANOVA from the package car^23^.

Cox Proportional-Hazards modelling was used to analyse survival. To do this, we used the “coxph” function from the package survival^24^.

Differences in survival between the control and a treatment group was analysed using a linear model (base R, “lm”) and post-hoc comparisons from the emmeans package^25^.

## Results

### Transient isoleucine deprivation increases nicotine tolerance of young and middle-aged flies

Previous research has shown that restricting dietary amino acids can confer stress resistance and benefit lifespan^26–30^. We have recently found that short term deprivation of a single essential amino acid can enhance nicotine tolerance of young adult flies, with seven-day exposure to an isoleucine dropout diet providing the greatest benefit^11^. However, we did not yet know if depriving a single amino acid in short bouts also extends lifespan and if so, when and for how long the restrictions need to be applied. To investigate this, we first assessed if flies could acquire nicotine tolerance from short bouts of amino acid deprivation as they aged. We maintained flies for one, two, three or five weeks (of their ∼9 week mean lifespan) on a nutritionally complete synthetic diet (Figure S1A), at which point we transferred them to a diet lacking isoleucine for 1, 3, 5 or 7 days (Figure S1.B), and then exposed the flies to a lethal dose of nicotine and measured their survival (Figure 1.A).

**Figure 1.**
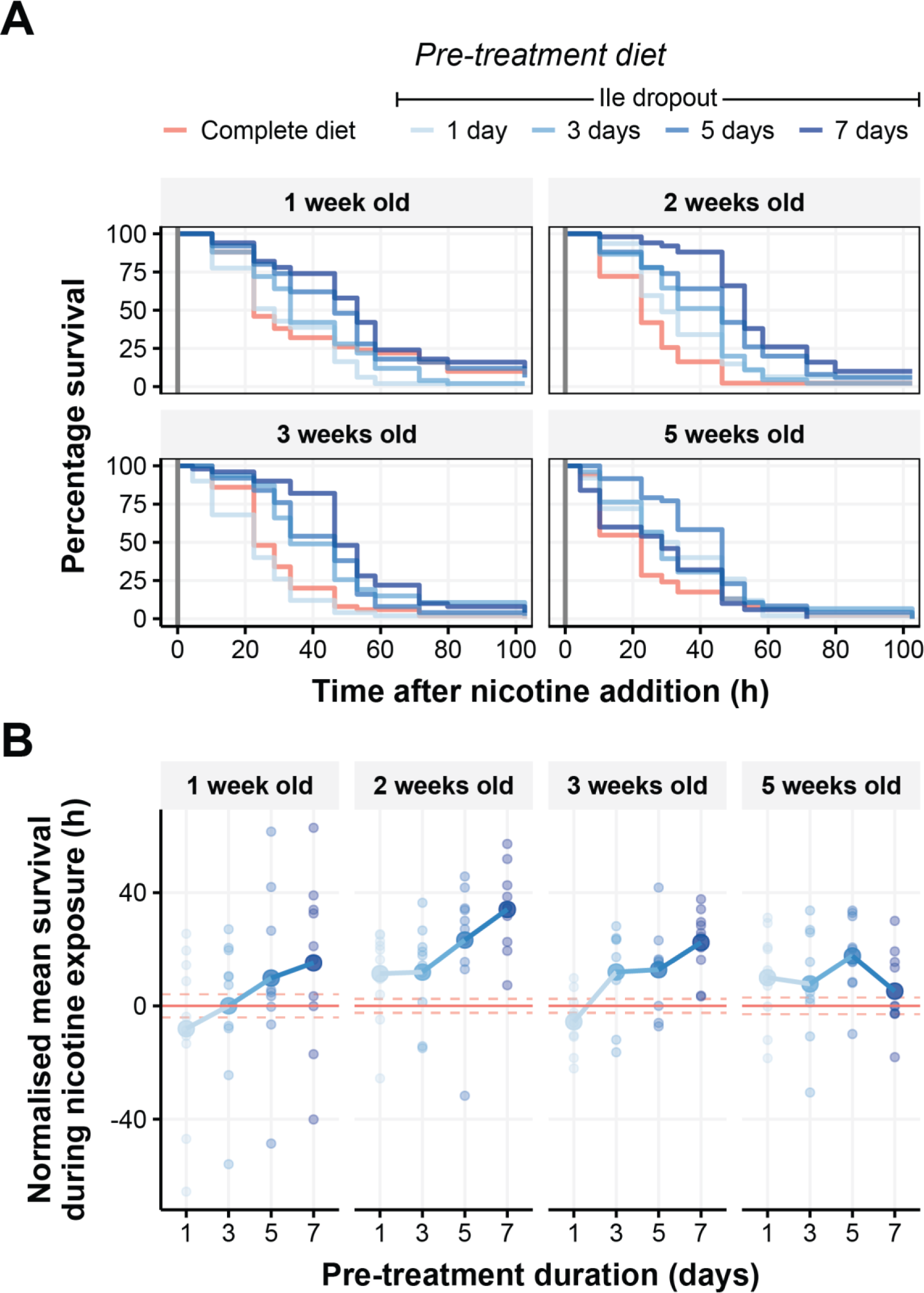
Short-term isoleucine deprivation confers nicotine resistance on young and middle-aged flies. Flies were pre-treated at either 1, 2, 3 or 5 weeks of age with an isoleucine dropout for either 1, 3, 5 or 7 days before chronic exposure to 0.83mg/mL of nicotine. (A) Survival curves of flies immediately following introduction of nicotine (indicated by vertical dark grey line). (B) Mean survival time of pre-treated flies when exposed to nicotine. The data are normalised to survival of flies fed a nutritionally complete synthetic diet (red horizontal line +/- SE indicated by dashed red lines), small circles represent mean lifespan for each replicate and large circles representing the group mean. Duration of pre-treatment positively correlated with survival upon nicotine exposure in all age groups except flies that were pre-treated at 5 weeks of age, where 5 days of pre-treatment was the most protective (Table S3). N = 50 flies per pre-treatment group for each age assayed.

Although flies were more susceptible to nicotine as they aged (Table S3, P < 0.01), increasing the time that flies were deprived of isoleucine also generally increased the flies’ survival during nicotine exposure (Table S4, P < 0.001). When separating the effects of isoleucine deprivation by age, we found that maximal protection was conferred by pre-treating flies with an isoleucine dropout for 7 days when they were 1, 2 or 3 weeks old, but when 5 weeks old, the most beneficial pre-treatment duration was 5 days (Figure 1B, Table S5). Thus, although diminished with age, flies still gain a protective benefit from short-term isoleucine deprivation at relatively advanced ages.

### Intermittent fasting of isoleucine only can extend lifespan

If resistance to a xenobiotic toxin, such as nicotine, reflects enhanced capacity to turnover life-threatening endobiotic compounds produced by metabolism^14^, we reasoned that short-term bouts of isoleucine deprivation may also confer longer life. We first selected the optimal duration of isoleucine deprivation using our previous data, and established that 7 days of isoleucine deprivation was ideal for 1- and 3-week-old flies and 5 days of deprivation was ideal for 5 week old flies. We then designed a range of dietary treatment plans in which flies were given either the nutritionally complete diet (control), or a schedule of isoleucine dropouts in all possible combinations of isoleucine deprivation on the first, third, and/or fifth weeks of adulthood (Figure 2A). At all other times, flies were provided with a nutritionally complete diet as we monitored their survival.

**Figure 2.**
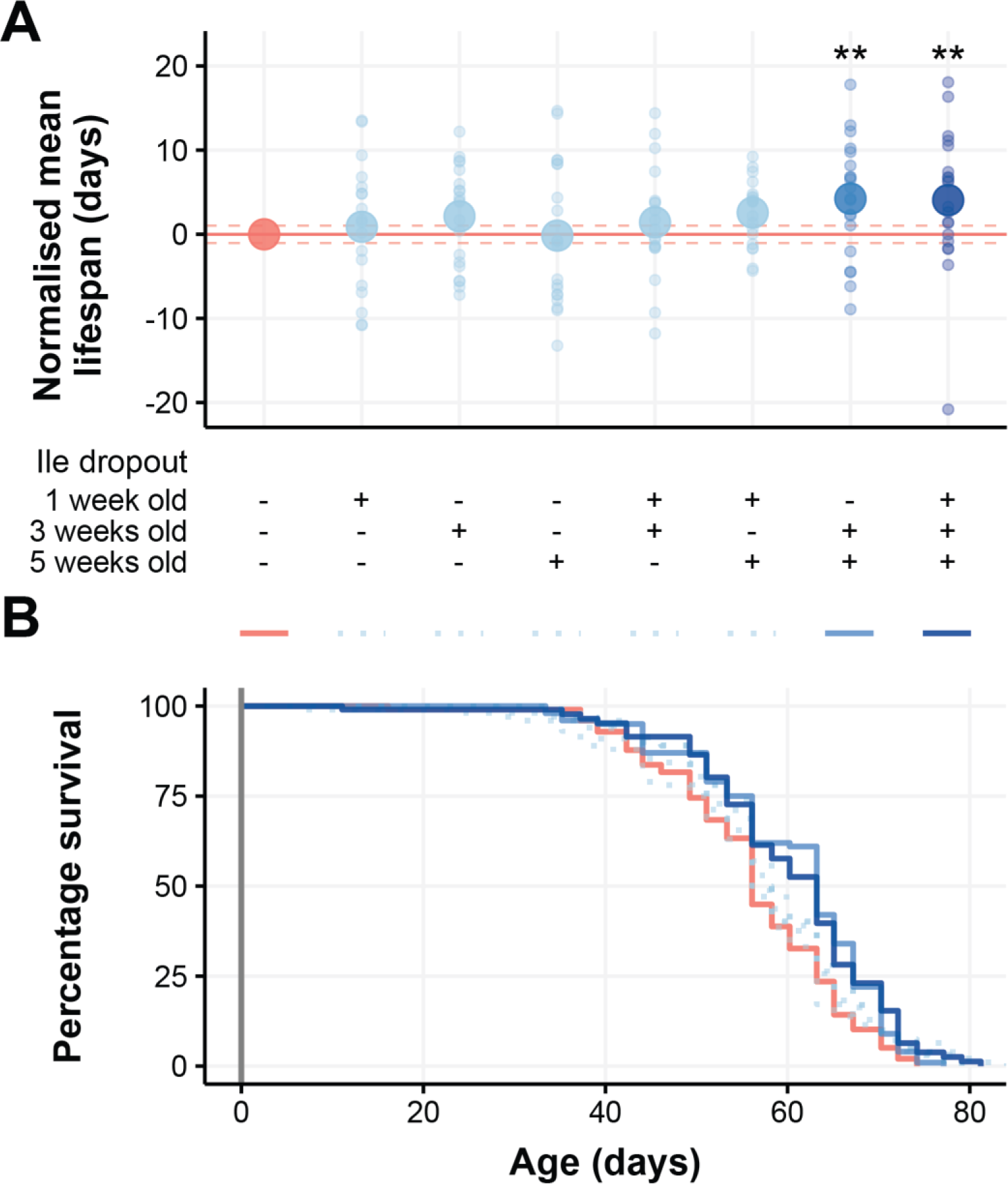
Transient periods of isoleucine restriction can extend lifespan when applied later in life. (A) Flies were assigned to a dietary regimen, which was comprised of a control cohort without Isoleucine deprivation, plus each combination of deprivation regime on 1, 3 and / or 5 weeks of adulthood. Isoleucine was omitted for 7 days at 1 week and 3 weeks old and for 5 days at 5 weeks old. The mean lifespan is represented as a large circle for the group, and small circles for each replicate. Flies that were deprived of isoleucine at both 3 and 5 weeks of age were significantly longer lived than the flies maintained on the nutritionally complete diet, irrespective of any prior isoleucine deprivation. (B) Survival curves over the duration of the flies’ lifespan, illustrating the extension conferred by removing isoleucine at both 3 and 5 weeks of age. N = 100 flies per dietary regimen.

We found that lifespan was significantly extended when the flies were deprived of isoleucine at 3 and 5 weeks of age, irrespective of whether or not they were also exposed to an isoleucine dropout diet at 1 week (Figure 2. A and B, Table S6). None of the other schedules of isoleucine dropout exposure affected lifespan when compared with control flies that were not exposed to isoleucine dropout food. These data suggest that the beneficial effects of isoleucine deprivation are cumulative, since more than one bout of treatment was required to extend life. They also indicate that the benefits of isoleucine deprivation extinguish over time, as the flies needed to be treated during the two later life windows to provide a benefit, while those exposed to any other pair of isoleucine deprivation treatments (weeks 1 & 3 or 1 & 5) were not long lived.

### Isoleucine deprivation only transiently protects against nicotine exposure

Our data are consistent with a model in which enhanced detoxification capacity protects flies against lifespan limiting damage. If so, our treatments could act either by clearing some life limiting damage during the time of isoleucine restriction, or by periodically boosting the activity of protective systems that is then sustained at a persistently higher level. To differentiate between these possibilities, we assayed flies for nicotine resistance either immediately after exposure to isoleucine deprivation (as in all previous assays) or using flies that had been returned to a complete diet for three days immediately following isoleucine deprivation. Interestingly, when flies were allowed to recover for three days on a complete diet after isoleucine deprivation, they completely lost the resistance to nicotine (Figure 3. Table S7). These data indicate that whatever the protective effect of intermittent isoleucine deprivation for lifespan, it is either brought about by transient clearing of damage during the bouts of isoleucine deprivation and / or by a mechanism that is unrelated to enhanced capacity to detoxify nicotine.

**Figure 3.**
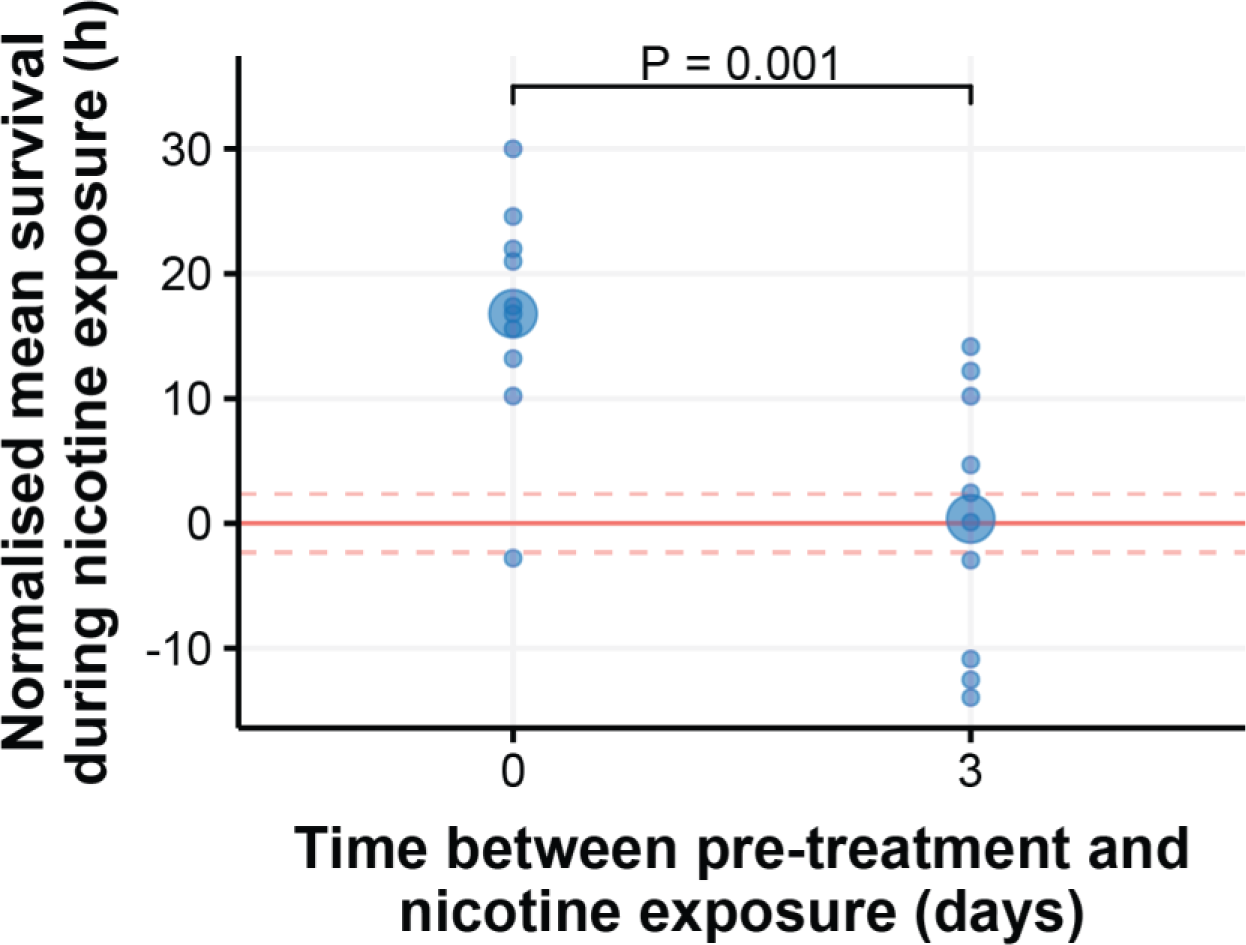
Removing isoleucine from the diet only protects flies against immediate exposure to nicotine. 1 week old flies were pre-treated with an isoleucine dropout for 7 days and then they were exposed to nicotine either immediately or after 3 days of being fed a complete diet. Flies that were immediately exposed to nicotine were protected by their pre-treatment, whereas flies exposed after three days of a complete diet did not benefit from pre-treatment. N = 50 flies per pre-treatment group and for each complete diet control.

## Discussion

In this study, we investigated the effects of short-term bouts of isoleucine deprivation on nicotine tolerance and lifespan in *Drosophila melanogaster*. We found that depriving flies of isoleucine conferred nicotine resistance at various ages and that the optimal duration of pre-treatment decreases as flies age. Strikingly, we observed that two bouts of transient isoleucine deprivation later in life was sufficient to extend lifespan regardless of whether the flies were also deprived of isoleucine early in adulthood. These results indicate a cumulative and time-dependent benefit of short-term isoleucine deprivation, as lifespan extension was only observed when flies experienced two bouts of isoleucine deprivation. However, these cumulative benefits are only present when treatments are close together and later in life, as flies weren’t longer lived when they were deprived of isoleucine at 1 and 5 weeks of age, or 1 and 3 weeks of age. Finally, our data suggest that intermittent isoleucine fasting is unlikely to extend lifespan by sustained activation of detoxification pathways, as flies lose their enhanced nicotine tolerance if they are maintained on a complete diet for three days in between isoleucine deprivation and nicotine poisoning. This suggests that some of the physiological consequences of isoleucine deprivation are not maintained when flies return to a complete diet, and that mechanisms extending lifespan may involve transient damage clearance during isoleucine deprivation or a mechanism unrelated to detoxification.

One possible explanation for our data is that our nutritionally complete synthetic diet contains too much isoleucine and removing it alleviates specific deleterious effects associated with isoleucine “poisoning”. However, the nutritionally complete diet we use was designed to optimise lifespan to a level equivalent to DR on a natural sugar-yeast medium^16 18^, suggesting that isoleucine fasting in this study does not simply alleviate isoleucine toxicity. It would be interesting to investigate whether the observed lifespan extension is isoleucine specific or if it can be replicated by short bouts of deprivation for another essential amino acid, such as lysine, which we previously found to be the second most potent in terms of nicotine resistance^11^. If intermittently depriving other essential amino acids can also increase lifespan beyond that of DR, this would point towards mechanisms underlying ageing that are not being fully targeted by current DR protocols, as the nutritionally complete synthetic diet mimics the lifespan under DR.

Intermittent isoleucine deprivation may activate the detoxification and / or antioxidant systems, both of which have been associated with longevity^15,31^. In long-lived insulin signalling mutants across different species, detoxification genes are strongly upregulated^32^. Similarly, long-lived insulin mutant flies are more resistant to DDT^33^, another neurotoxic insecticide that is eliminated by the detoxification system^34^. However, both in these previous studies and our current work, enhanced detoxification can be separated from longevity, suggesting the involvement of other protective mechanisms. Genetic suppression of the antioxidants superoxide dismutase or catalase severely compromises lifespan of flies, and similarly, pharmacologically inhibition of superoxide dismutase reduces the lifespan of long-lived flies but not flies with a “normal” lifespan^35^. Conversely, overexpressing these antioxidants can extend lifespan^36^. It is possible that isoleucine deprivation upregulates both detoxification and antioxidant pathways. It would be interesting to transiently upregulate these processes genetically or pharmacologically on the same schedule as isoleucine deprivation to see if they increase lifespan, or if suppressing these pathways during isoleucine deprivation blocks the extension of lifespan. Further, investigating the overlapping molecular signatures between isoleucine deprivation, reduced insulin signalling, and enhanced detoxification and antioxidant pathways could provide valuable insights into the other protective functions that are enhanced by amino acid deprivation.

Our data are also consistent with a model in which endogenous damage is cleared during bouts of isoleucine deprivation, and these effects accumulate to generate beneficial outcomes on lifespan. A reasonable mechanistic explanation for this is that isoleucine deprivation activates the integrated stress response (ISR) through the intracellular amino acid sensor and kinase GCN2^37^. This reduces global translation through phosphorylating eIF2α (eukaryotic translation Initiation Factor-2α), which also induces autophagy through suppression of mTORC1^38^. Together, these processes should transiently induce autophagy to enhance proteostasis that may suppress ageing, similarly to the hormetic effects of heat shock in *C. elegans*^39^. It would be interesting to see if transient activation of the ISR and/or suppression of mTORC1 by other means during weeks 3 and 5 of adulthood could also extend the lifespan of flies. Further, characterising the molecular signatures associated with temporary isoleucine deprivation, as well as the connection between nutrient signalling and lifespan, should provide evidence for these effects or reveal new protective mechanisms involved in longevity. Future studies should also investigate the nature of the possible damage accumulating in ageing flies that may be cleared by isoleucine deprivation. If done in a tissue-specific fashion, it would present an opportunity for short-term tissue-specific manipulations of nutrient sensing pathways to improve lifespan.

Interestingly, we found that bouts of isoleucine fasting were only beneficial when implemented later in life. This finding directly contrasts other dietary manipulations, which demonstrate that intermittent fasting^40^ or transient methionine restriction^41^ extend fly lifespan when implemented early, but not later, in life. Similarly, fly lifespan can be extended by rapamycin treatment in the first 30 days of life, but the benefit declines when they are treated later in life^42^. One important difference between protocols could be the schedule of treatments: these were longer (2-4 weeks) in previous studies and only a single bout rather than the shorter, repeated bouts that we implemented. It would be interesting to see if these other dietary manipulations could extend lifespan on the same schedule as isoleucine deprivation. In particular, we predict that short bouts of rapamycin treatment during weeks 3 and 5 of adulthood would extend lifespan, as we have shown that rapamycin phenocopies isoleucine deprivation in protection against nicotine^11^. Similarly, it would be interesting to see if depriving young flies of isoleucine for a longer time could extend lifespan, as early rapamycin treatment and methionine restriction do. These experiments are important to establish applications that promote longevity when implemented in older individuals.

## Conclusions

Our study presents exciting findings that repeated short bouts of single amino acid deprivation can enhance stress resistance and lifespan. This presents a promising avenue for greater precision in dietary interventions, applied later in life, to promote healthy ageing. Future studies to elucidate the mechanistic basis of this finding could pave the way for pharmacological approaches to mimic these benefits, thus achieving geroprotection from shorter periods of less-invasive interventions.

## Acknowledgements

The Project was supported by a Longevity Impetus Grant from Norn Group.

## Supplementary materials

**Supplementary Table 1.**
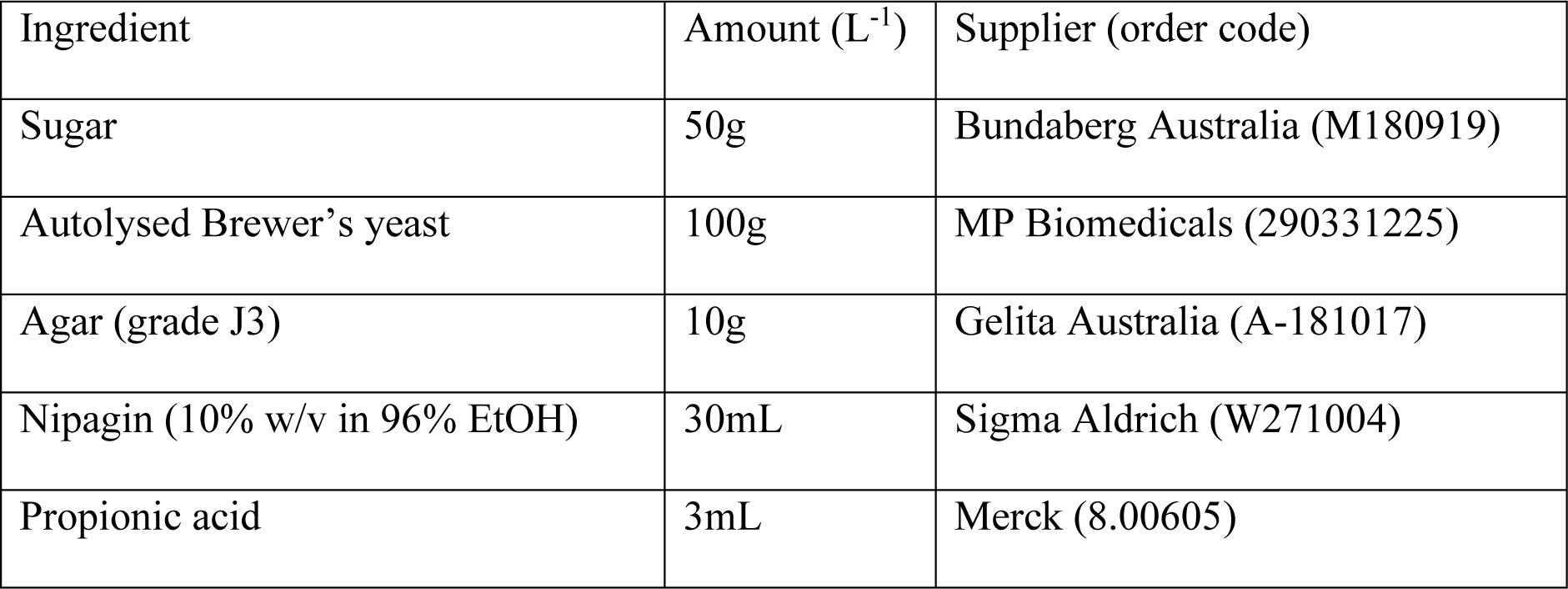
Ingredients in 1L of Sugar Yeast medium.

**Supplementary Table 2.**
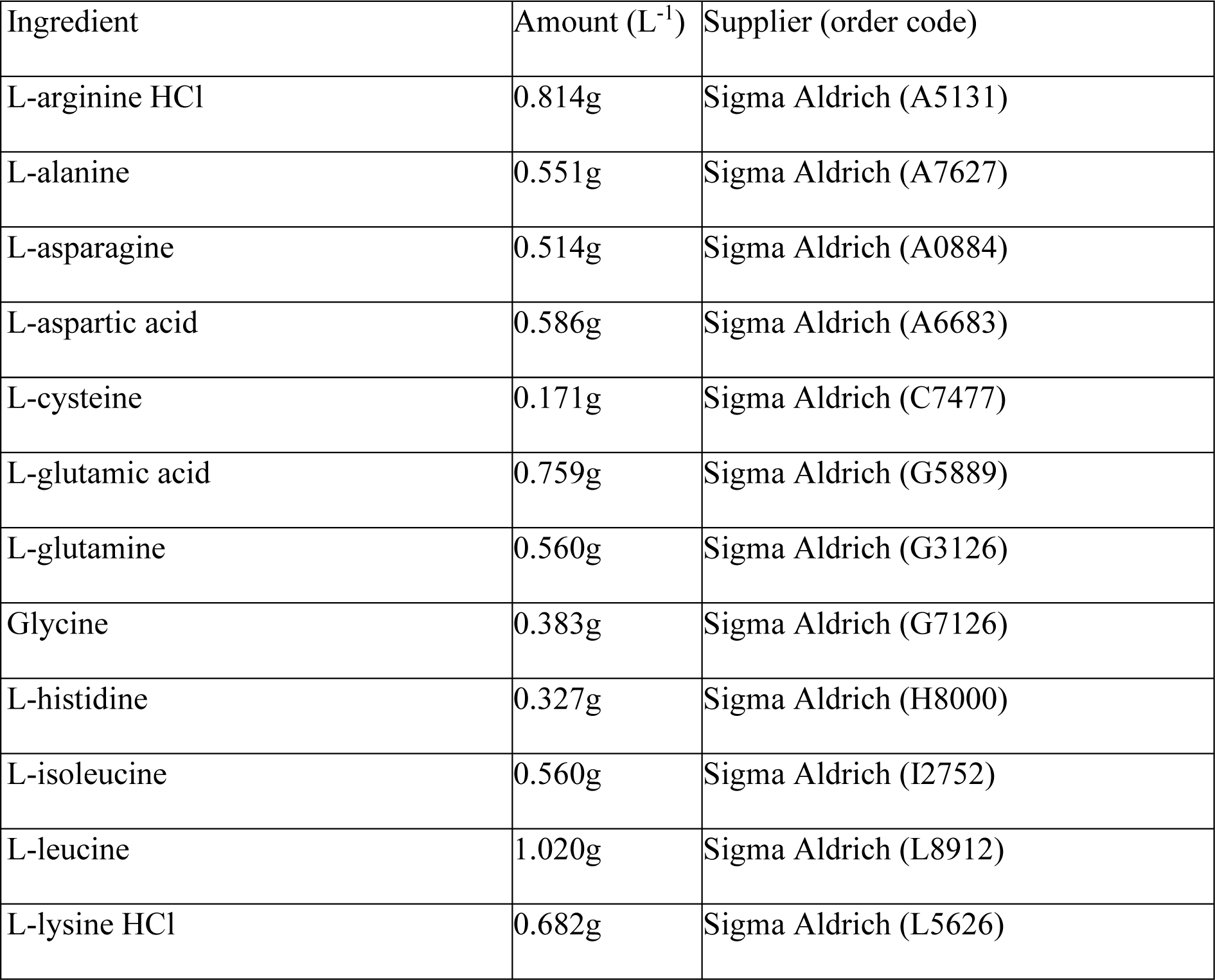
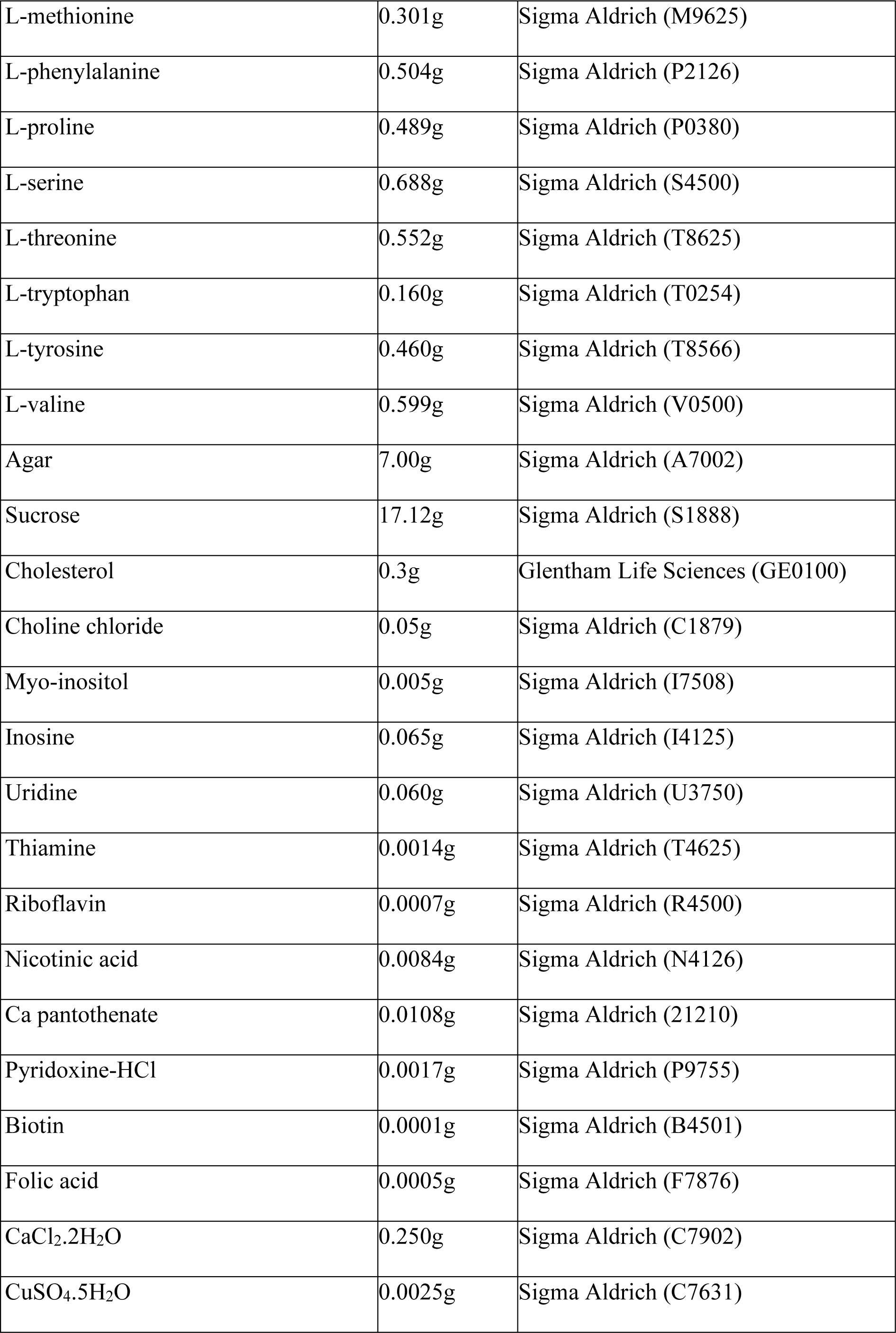
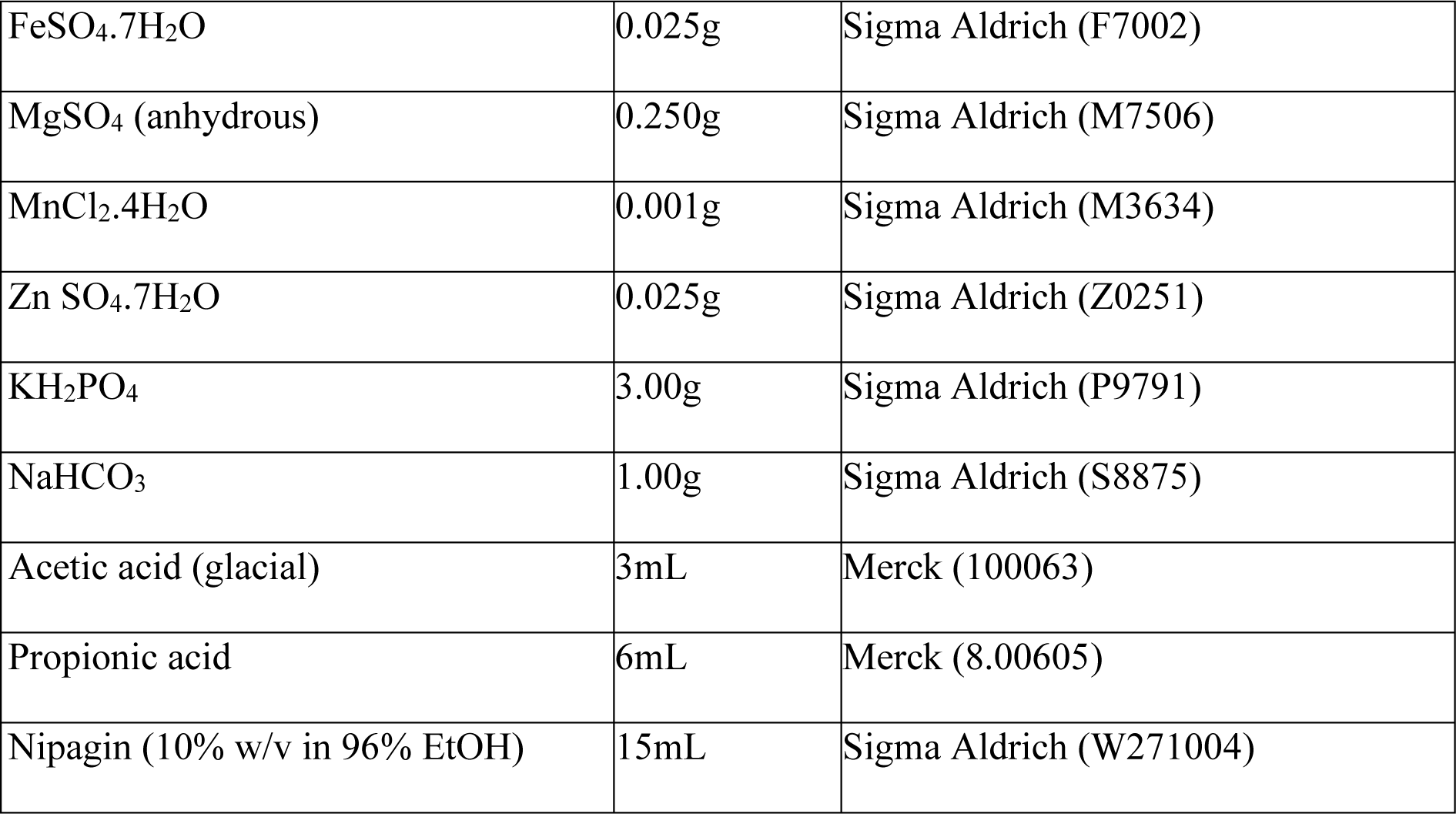
Ingredients in 1L of complete synthetic medium.

| Ingredient | Amount (L<sup>-1</sup>) | Supplier (order code) |
| --- | --- | --- |
| L-arginine HCl | 0.814g | Sigma Aldrich (A5131) |
| L-alanine | 0.551g | Sigma Aldrich (A7627) |
| L-asparagine | 0.514g | Sigma Aldrich (A0884) |
| L-aspartic acid | 0.586g | Sigma Aldrich (A6683) |
| L-cysteine | 0.171g | Sigma Aldrich (C7477) |
| L-glutamic acid | 0.759g | Sigma Aldrich (G5889) |
| L-glutamine | 0.560g | Sigma Aldrich (G3126) |
| Glycine | 0.383g | Sigma Aldrich (G7126) |
| L-histidine | 0.327g | Sigma Aldrich (H8000) |
| L-isoleucine | 0.560g | Sigma Aldrich (I2752) |
| L-leucine | 1.020g | Sigma Aldrich (L8912) |
| L-lysine HCl | 0.682g | Sigma Aldrich (L5626) |
| L-methionine | 0.301g | Sigma Aldrich (M9625) |
| L-phenylalanine | 0.504g | Sigma Aldrich (P2126) |
| L-proline | 0.489g | Sigma Aldrich (P0380) |
| L-serine | 0.688g | Sigma Aldrich (S4500) |
| L-threonine | 0.552g | Sigma Aldrich (T8625) |
| L-tryptophan | 0.160g | Sigma Aldrich (T0254) |
| L-tyrosine | 0.460g | Sigma Aldrich (T8566) |
| L-valine | 0.599g | Sigma Aldrich (V0500) |
| Agar | 7.00g | Sigma Aldrich (A7002) |
| Sucrose | 17.12g | Sigma Aldrich (S1888) |
| Cholesterol | 0.3g | Glentham Life Sciences (GE0100) |
| Choline chloride | 0.05g | Sigma Aldrich (C1879) |
| Myo-inositol | 0.005g | Sigma Aldrich (I7508) |
| Inosine | 0.065g | Sigma Aldrich (I4125) |
| Uridine | 0.060g | Sigma Aldrich (U3750) |
| Thiamine | 0.0014g | Sigma Aldrich (T4625) |
| Riboflavin | 0.0007g | Sigma Aldrich (R4500) |
| Nicotinic acid | 0.0084g | Sigma Aldrich (N4126) |
| Ca pantothenate | 0.0108g | Sigma Aldrich (21210) |
| Pyridoxine-HCl | 0.0017g | Sigma Aldrich (P9755) |
| Biotin | 0.0001g | Sigma Aldrich (B4501) |
| Folic acid | 0.0005g | Sigma Aldrich (F7876) |
| CaCl<sub>2</sub>.2H<sub>2</sub>O | 0.250g | Sigma Aldrich (C7902) |
| CuSO<sub>4</sub>.5H<sub>2</sub>O | 0.0025g | Sigma Aldrich (C7631) |
| FeSO<sub>4</sub>·7H<sub>2</sub>O | 0.025g | Sigma Aldrich (F7002) |
| MgSO<sub>4</sub> (anhydrous) | 0.250g | Sigma Aldrich (M7506) |
| MnCl<sub>2</sub>·4H<sub>2</sub>O | 0.001g | Sigma Aldrich (M3634) |
| Zn SO<sub>4</sub>·7H<sub>2</sub>O | 0.025g | Sigma Aldrich (Z0251) |
| KH<sub>2</sub>PO<sub>4</sub> | 3.00g | Sigma Aldrich (P9791) |
| NaHCO<sub>3</sub> | 1.00g | Sigma Aldrich (S8875) |
| Acetic acid (glacial) | 3mL | Merck (100063) |
| Propionic acid | 6mL | Merck (8.00605) |
| Nipagin (10% w/v in 96% EtOH) | 15mL | Sigma Aldrich (W271004) |

**Supplementary Table 3.** Survival model of flies that were fed a complete diet during weeks 1, 2, 3, or 5 of age and then exposed to nicotine. Summary of cox-proportional hazards modelling with age as a continuous variable. Confidence level = 95%.

| Terms | Estimate | Hazard ratio | SE | Z value |
| --- | --- | --- | --- | --- |
| Exposure Age | 0.14 | 1.15 | 0.05 | **2.83\*\*** |
| ***p < 0.001, **p < 0.01, *p < 0.05 |  |  |  |  |

**Supplementary Table 4.**
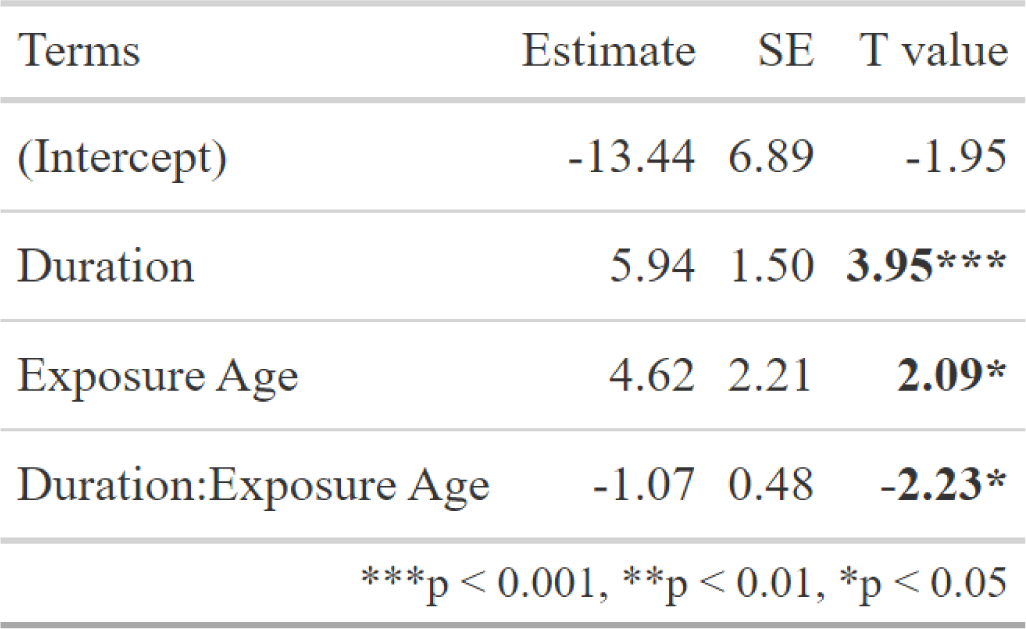
Effects of duration of pre-treatment and age on the difference in survival between pre-treated flies and control flies. Summary of the linear model that best represented the relationship with duration of pre-treatment and age as continuous variables. Confidence level = 95%.

| Terms | Estimate | SE | T value |
| --- | --- | --- | --- |
| (Intercept) | -13.44 | 6.89 | -1.95 |
| Duration | 5.94 | 1.50 | **3.95\*\*\*** |
| Exposure Age | 4.62 | 2.21 | **2.09\*** |
| Duration:Exposure Age | -1.07 | 0.48 | **-2.23\*** |
| ***p < 0.001, **p < 0.01, *p < 0.05 |  |  |  |

**Supplementary Table 5.**
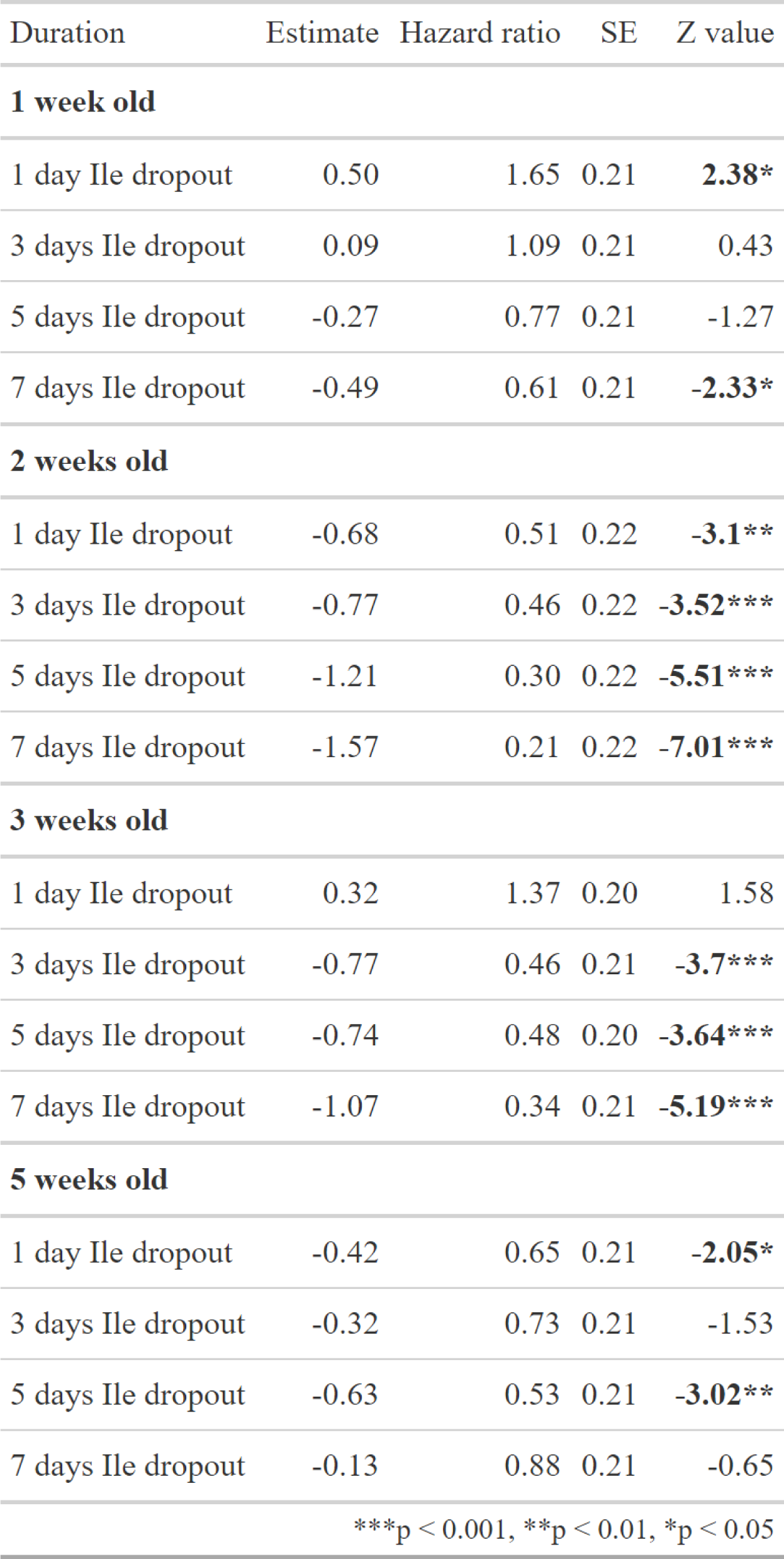
Differences in survival between flies that were pre-treated with an isoleucine dropout for 1, 3, 5 or 7 days compared to flies that were fed a complete diet, separated by age when pre-treated. Summary of cox-proportional hazards modelling with duration of pre-treatment and age as discrete variables. Confidence level = 95%Supplementary Table 6. Differences in survival between flies that were intermittently fasted of isoleucine and flies that were fed a complete diet for their lifespan. Summary of cox-proportional hazards modelling. The Terms column represents the sequence of Ile-deprivation treatments, where first is at 1 week of age, second at 3 weeks of age, and third at 5 weeks of age. Confidence level = 95%.

| Duration | Estimate | Hazard ratio | SE | Z value |
| --- | --- | --- | --- | --- |
| **1 week old** |  |  |  |  |
| 1 day Ile dropout | 0.50 | 1.65 | 0.21 | **2.38\*** |
| 3 days Ile dropout | 0.09 | 1.09 | 0.21 | 0.43 |
| 5 days Ile dropout | -0.27 | 0.77 | 0.21 | -1.27 |
| 7 days Ile dropout | -0.49 | 0.61 | 0.21 | **-2.33\*** |
| **2 weeks old** |  |  |  |  |
| 1 day Ile dropout | -0.68 | 0.51 | 0.22 | **-3.1\*\*** |
| 3 days Ile dropout | -0.77 | 0.46 | 0.22 | **-3.52\*\*\*** |
| 5 days Ile dropout | -1.21 | 0.30 | 0.22 | **-5.51\*\*\*** |
| 7 days Ile dropout | -1.57 | 0.21 | 0.22 | **-7.01\*\*\*** |
| **3 weeks old** |  |  |  |  |
| 1 day Ile dropout | 0.32 | 1.37 | 0.20 | 1.58 |
| 3 days Ile dropout | -0.77 | 0.46 | 0.21 | **-3.7\*\*\*** |
| 5 days Ile dropout | -0.74 | 0.48 | 0.20 | **-3.64\*\*\*** |
| 7 days Ile dropout | -1.07 | 0.34 | 0.21 | **-5.19\*\*\*** |
| **5 weeks old** |  |  |  |  |
| 1 day Ile dropout | -0.42 | 0.65 | 0.21 | **-2.05\*** |
| 3 days Ile dropout | -0.32 | 0.73 | 0.21 | -1.53 |
| 5 days Ile dropout | -0.63 | 0.53 | 0.21 | **-3.02\*\*** |
| 7 days Ile dropout | -0.13 | 0.88 | 0.21 | -0.65 |
| ***p < 0.001, **p < 0.01, *p < 0.05 |  |  |  |  |

**Supplementary Table 6.** Differences in survival between flies that were intermittently fasted of isoleucine and flies that were fed a complete diet for their lifespan. Summary of cox-proportional hazards modelling. The Terms column represents the sequence of Ile-deprivation treatments, where first is at 1 week of age, second at 3 weeks of age, and third at 5 weeks of age. Confidence level = 95%.

| Terms | Estimate | Hazard ratio | SE | Z value |
| --- | --- | --- | --- | --- |
| + - - : First only | -0.11 | 0.89 | 0.15 | -0.76 |
| - + - : Second only | -0.26 | 0.77 | 0.14 | -1.85 |
| - - + : Third only | -0.13 | 0.88 | 0.14 | -0.93 |
| + + - : First and second | -0.25 | 0.78 | 0.15 | -1.7 |
| + - + : First and third | -0.25 | 0.78 | 0.14 | -1.75 |
| - + + : Second and third | -0.39 | 0.68 | 0.14 | **-2.75\*\*** |
| + + + : All three | -0.44 | 0.64 | 0.15 | **-2.92\*\*** |
| ***p < 0.001, **p < 0.01, *p < 0.05 |  |  |  |  |

**Supplementary Table 7.** The effect of returning isoleucine to the flies’ diet for 3 days before nicotine exposure on the difference in survival between pre-treated flies and control flies. The linear model was tested using emmeans25 to determine whether the difference in survival was different from the control (where control = 0 days). Confidence level = 95%.

| Terms | Estimate | SE | DF | T Ratio |
| --- | --- | --- | --- | --- |
| 0 days | 16.80 | 3.06 | 18 | **5.5\*\*\*** |
| 3 days | 0.35 | 3.06 | 18 | 0.11 |

\*\*\*p < 0.001, \*\*p < 0.01, \*p < 0.05

**Supplementary Figure 1.**
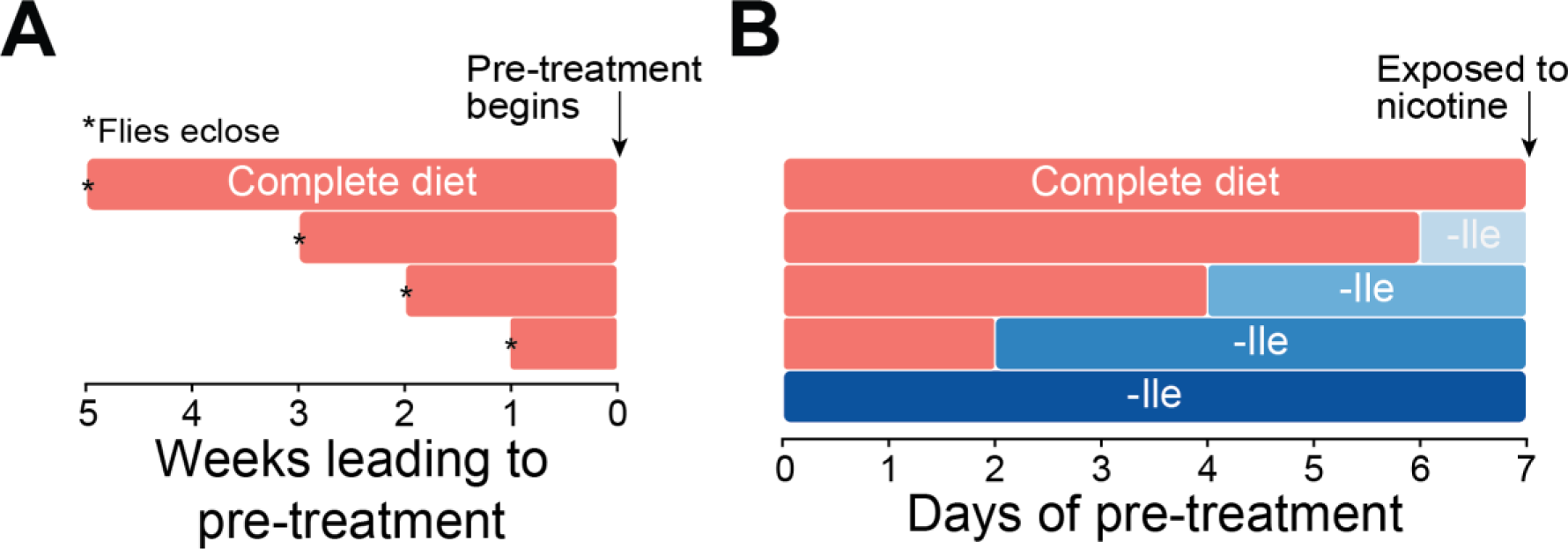
Methods depicting the treatment timeline for data shown in Figure 1. **(A)** Controlled density egg lays were timed to ensure that flies eclosed at 5, 3, 2 and 1 week prior to the Ile-deprivation pre-treatment period. Flies were maintained on a complete, synthetic diet during this maintenance period. (**B**) During the pre-treatment period, flies were placed onto an isoleucine dropout for 7, 5, 3, 1 or 0 days immediately prior to nicotine exposure. Nicotine treatment was applied to all groups contemporaneously.

